# Phase of transcranial alternating current stimulation modulates working memory processing speed

**DOI:** 10.64898/2026.05.26.727793

**Authors:** Julie Dimmendaal, Xiaojuan Wang, Bas J. Dijkslag, Laura E. Huizinga, Sander Maalderink, Marijn Priest, Femke J.E. van Dam, Mark M. Span, Miles Wischnewski

**Affiliations:** Department of Psychology, University of Groningen, Groningen, The Netherlands; Medical School, Tianjin University, Tianjin, China; Department of Cognitive Neuroscience, Maastricht University, Maastricht, The Netherlands

**Keywords:** Working memory, Transcranial alternating current stimulation, Oscillation phase, Inter-individual variability

## Abstract

**Background:** Theta-frequency transcranial alternating current stimulation (tACS) over prefrontal cortex has been proposed to modulate working memory (WM), yet behavioral effects are often inconsistent. One potential source of variability is the tACS phase during stimulus presentation.

**Objective:** We tested whether behavioral performance during WM depends on the phase of prefrontal theta-tACS.

**Methods:** Twenty participants completed two sessions of prefrontal 4 Hz tACS in a within-subject design, receiving active and sham stimulation in separate sessions. Participants performed a visuospatial change detection task (CDT) and a verbal N-back task. Stimulation effects on overall accuracy and reaction time were analyzed. Subsequently, phase-specific analyses related stimulation phase at task-relevant stimulus presentation to behavioral performance using circular regression models. Preferred phases across participants were tested using Rayleigh tests.

**Results:** No significant overall effects of active compared with sham tACS on accuracy or reaction time were observed in either task. However, phase-specific analyses revealed stronger phase-dependent modulation of reaction time during active tACS compared with sham. In the CDT, this effect was present across difficulty levels, whereas in the N-back task it was observed only in the 3-back condition. No reliable phase-dependent effects were observed for accuracy. Preferred phases varied across participants and did not cluster around a common phase.

**Conclusions:** Prefrontal theta-tACS can modulate WM performance in a phase-dependent manner even in the absence of average behavioral effects. The observation of phase-dependent reaction-time modulation across visuospatial and verbal WM tasks suggests that stimulation phase may be a relevant source of variability across cognitive domains.

## 1. Introduction

Working memory (WM) enables the temporary maintenance and manipulation of limited information [1–3]. Substantial evidence links WM maintenance to activity in the prefrontal cortex [4–8]. While various studies demonstrated persistently elevated activity during WM maintenance [9–11], others have challenged this notion [6,12,13]. Rather, WM items may be processed and recalled through rhythmic activity [3,6,14]. Moreover, invasive recordings in macaques and surgical patients have shown that neural oscillation phase determines recall performance in WM [15–19]. However, evidence on the importance of oscillation phase in WM in the healthy human brain is currently limited.

Transcranial alternating current stimulation (tACS) can be used to study the causal role of neural oscillations [20–23]. Indeed, prefrontal theta-frequency tACS has been shown to improve WM performance [24–29]. A recent meta-analysis of 28 placebo-controlled tACS studies demonstrated that prefrontal theta tACS improves WM performance with a small-to-moderate effect size [30]. However, effects remain inconsistent across studies, with considerable inter-study, inter-individual, and intra-individual variability [30,31]. For instance, tACS effects on WM may depend on task difficulty and participants’ baseline performance [29,32–34]. Debnath et al. [29] reported that theta tACS over the left DLPFC reduced reaction times during WM tasks only under high-load conditions. Furthermore, stimulation parameters, particularly the exact frequency, can also affect tACS efficacy [24,35]. Aktürk et al. [24] found that theta-frequency tACS applied at a frequency slightly below an individual’s endogenous theta frequency improved memory performance compared to stimulation delivered above the individual theta frequency and sham stimulation.

One potential, often overlooked, source of variability is the phase of the tACS oscillation. Since tACS can entrain neuronal firing to specific phases of the applied sinusoidal current, it modulates the timing of neural activity [36–39]. Thus, tACS effects may depend not only on stimulation location, intensity, and frequency, but also on stimulation phase [40–47]. For instance, motor evoked responses following transcranial magnetic stimulation are modulated by the phase of the applied tACS oscillation [40,41,45]. Furthermore, Fiene and colleagues demonstrated that the amplitude of visually evoked steady state responses, as well as the brightness perception of a visual stimulus, depend on the phase of tACS [42,43]. Similar tACS phase-dependent sensory modulation was observed in the auditory domain, with auditory detection performance varying across the tACS cycle [46].

Given that WM performance may depend on the prefrontal theta phase [15,16], effects of tACS may be affected by the exact timing of the presented stimuli relative to the tACS phase. In the present study, the relation between tACS phase and performance in a spatial and verbal WM task was assessed. Specifically, we investigated whether the tACS phase at which WM stimuli are shown affects task accuracy and reaction time (RT). Furthermore, we investigated whether any phase dependencies are consistent across participants.

## 2. Methods

### 2.1. Participants

A total of 25 healthy adults were recruited for this study, with the final analyzed sample consisting of N=20 (age range 18-35 years). Two recruited participants were excluded from the analyses because they performed a task with stimulus durations different from those used in the analyses. Three participants were excluded because they participated in only a single session. Exclusion criteria comprised the presence of metal implants in or near the head, a history of epilepsy, dermatological conditions near the site of stimulation, active psychiatric disorders, pregnancy, and color blindness [48]. The study was approved by the ethics board of the Faculty of Behavioural and Social Sciences at the University of Groningen. Participants provided written informed consent.

### 2.2. Design

This study used a placebo-controlled, within-subjects design. Each participant completed two experimental sessions at least one week apart to limit carry-over effects, with both sessions scheduled at the same time of day to limit circadian effects on task performance [49]. Participants received active theta tACS in one session and sham stimulation in the other. In each session, they performed a change detection task (CDT) with three difficulty levels and an N-back task with two difficulty levels. Session order was randomized and counterbalanced, and participants were blind to the stimulation condition. Each session lasted approximately 75 minutes.

An a priori sample size calculation was performed using G*Power 3.1 [50]. The power analysis assumed the CDT as the main task and a repeated measures ANOVA as the statistical test. A medium effect size of Cohen’s f = 0.25, power of 1-β = 0.9, significance level of α = 0.05, and correlation between measurements of r = 0.75 [51,52] were assumed. This resulted in a suggested sample size of N=19. As such, the realized power with our final sample of N=20 was 1-β = 0.925 for the CDT and 1-β = 0.851 for the N-back task.

### 2.3. Working memory tasks

A CDT [34,35,53,54] and an N-back task [34,55–57] were used to test visuospatial and verbal WM, respectively. The tasks were programmed and run using OpenSesame software (v4.1) [58]. Both tasks were presented on a 22-inch monitor with a screen resolution of 1920×1080 pixels, with participants being seated approximately 80 cm from the monitor.

#### 2.3.1. CDT

For the CDT, an array of colored squares was presented at random locations on the screen. After a delay period, participants indicated whether the color of one of the squares had changed (Figure 1A). Task difficulty was manipulated by varying the number of squares on the screen across three levels: moderate (four squares), hard (six squares), and very hard (eight squares). Each square (50 × 50 pixels) was displayed in one of 21 implemented colors and presented at quasi-random screen locations. To ensure an even spatial distribution, the screen was divided into four quadrants, and squares were presented at random, non-overlapping positions within these quadrants.

**Figure 1.**
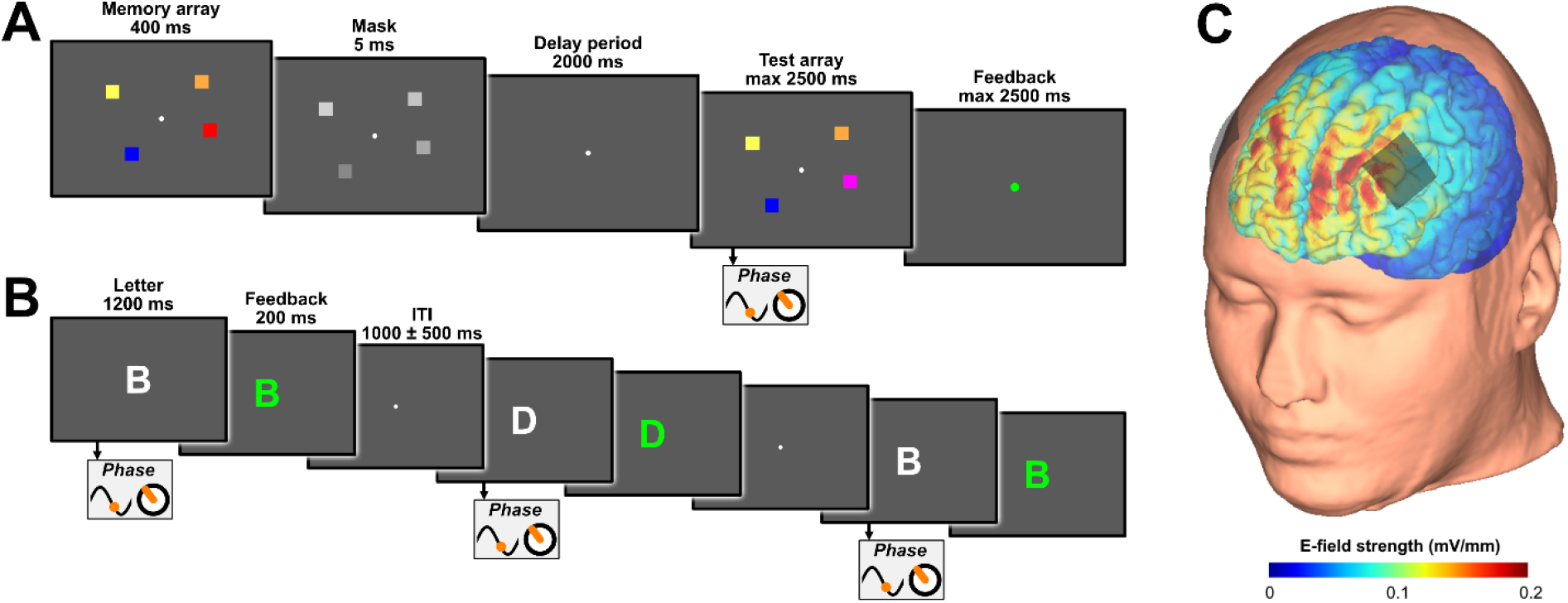
Experimental tasks and stimulation montage. (A) Example trial sequence of the change detection task (CDT). Participants viewed a memory array, followed by a brief mask, a delay period, and a test array, and indicated whether one of the squares had changed color. (B) Example sequence (three trials) of the N-back task. Participants responded to target letters depending on whether the current letter matched the one presented N trials earlier. Phase symbols indicate stimulus events at which the tACS phase was extracted for analysis. (C) Electric field model of the prefrontal tACS montage, showing the estimated field strength over the cortex.

After a fixation cross (200 ms), the first array of squares (memory array) was presented for 400 ms. Subsequently, a mask was shown for 5 ms, during which the color of all squares changed. Participants were instructed to ignore the mask. The mask was included to increase WM demands during the delay period. The mask was followed by a 2000 ms delay period during which only a fixation cross was displayed. Next, the second array of squares (test array) was presented. This array was either identical to the memory array or contained a color change in one square. The test array remained on screen until the participant responded or for a maximum of 2500 ms. Participants responded by pressing one of two buttons to indicate whether a change had occurred or not. Change and no-change trials were presented in a 1:1 ratio. Finally, a green or red dot, shown for 300 ms indicated correct or incorrect responses.

In total, the CDT consisted of 180 trials divided into six blocks of 30 trials. Block difficulty followed the same fixed sequence for all participants: moderate, hard, very hard, repeated twice. Before the main task, participants completed a practice run without tACS consisting of 15 trials at both the moderate and hard difficulty levels.

#### 2.3.2. N-back task

For the N-back task, participants indicated whether the currently presented letter matched the one shown N trials earlier (Figure 1B). Two difficulty levels were included: 2-back and 3-back. After a fixation cross (200 ms), a stimulus was presented (32 pixels in height), consisting of one of the first eight letters (A–H) of the Latin alphabet. Each letter remained on screen for a maximum of 1200 ms, during which participants could respond. Participants were instructed to press a button when the presented letter was a match and to withhold a response when it was a non-match. Feedback was provided after each trial by changing the letter color to green (correct) or red (incorrect) for 200 ms. Between trials, a jittered blank-screen interval of 500–1500 ms was used.

In total, the task consisted of 240 trials, with 120 trials each for the 2-back and 3-back conditions. Before the main task, participants completed a practice run without tACS consisting of 40 trials for both the 2-back and 3-back conditions.

### 2.4 tACS

TACS was applied using a NeuroConn DC-stimulator Plus (NeuroConn GmbH, Ilmenau). During the experiment, active tACS was applied at 4 Hz with an intensity of 1.5 mA (peak-to-peak) for 30 minutes. Rubber electrodes were used together with conductive paste (Ten20 Conductive Paste) placed over the bilateral prefrontal cortex (F3 and F4 according to the international 10-20 system). An estimation of the electric field distribution was done using SimNIBS [59], and is shown in Figure 1C. A ramp up/down of 30 seconds was used at the beginning and end of stimulation. The impedance was kept below 10 kΩ throughout the experiment. During sham stimulation, the same intensity, montage, and frequency were used, but stimulation was applied for 90 seconds (30 seconds ramp-up, 30 seconds stimulation, 30 seconds ramp-down).

Phase was monitored using an induction ring placed around the cable connected to the F3 electrode. For sham stimulation, no signal was recorded after the initial ramp-up/down period, and a synthetic 4 Hz sinusoidal function was used to assign stimulus onset to phase.

### 2.5 Data processing and analysis

#### 2.5.1. Phase quantification and analysis

To determine the stimulation phase at stimulus onset, the recorded tACS signal was band-pass filtered between 3 and 5 Hz using a fourth-order zero-phase Butterworth filter centered at the stimulation frequency of 4 Hz. The instantaneous phase was derived from the filtered signal using the Hilbert transform, yielding phase angles in radians. Task markers provided through OpenSesame were used to extract the corresponding phase at stimulus onset. For the CDT, the phase was determined at the presentation of the test array. For the N-back task, the phase was determined at the presentation of each letter.

To assess phase–behavior associations (for both CDT and N-back), circular regression analyses were used at the participant level, using the CircStat toolbox [60] implemented in Matlab (version 2024b). Regression models were calculated for each difficulty level, and the same approach was used for the CDT and N-back tasks.

The relation between RT and phase was computed via circular-linear regression models, with the sine and cosine of phase as predictors.

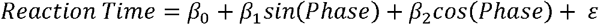

Regression coefficients, R^2^, circular-linear correlation (r_circ_), model significance, modulation amplitude, and preferred phase were extracted. For the CDT, all trials were considered in the regression. For the N-back task, because participants responded only when a match occurred, only trials with a response were used.

The relation between accuracy and phase was computed using circular and binary logistic regression models using the sine and cosine of phase as predictors.

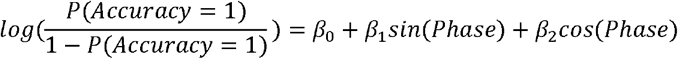

Since standard R^2^ is not defined for logistic regression, we report a pseudo-R^2^ that reflects the improvement of the fitted model over a null model. In addition, regression coefficients, model significance, modulation amplitude, and preferred phase were extracted. For comparability with the r_circ_ coefficients obtained for RT, the square root of the deviance-based pseudo-R^2^ from the logistic regression models was used as a pseudo-correlation index for accuracy (pseudo-r_circ_).

To investigate phase–behavior relationships between participants, they were plotted on a polar scatterplot where the angle refers to the preferred phase and distance from the center reflects the Fisher z-transformed (pseudo-)r_circ_ values. For accuracy, pseudo-r values derived from pseudo-R^2^ were used as an analogous effect-size measure to facilitate comparability with the RT analyses. Preferred phase reflects the phase angle at which the modeled effect was maximal and was calculated from the sine and cosine coefficients as:

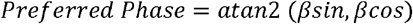

The preferred phase corresponds to the maximum value of the regression fit. For accuracy, this reflects the phase with the highest probability of being correct. For RT, because lower values indicate better performance, the calculated preferred phase was shifted by 180° to represent the phase associated with the fastest predicted responses. Accordingly, when referring to the preferred phase for RT, we refer to the phase with the fastest predicted responses.

#### 2.5.2. Statistical analysis

Statistical analyses were conducted using JASP (version 0.18) and MATLAB (version 2024b). To assess general behavioral effects, repeated-measures analyses of variance (RM-ANOVAs) were performed with accuracy and RT as dependent variables. For the CDT, a 3 × 2 RM-ANOVA was used with Difficulty (moderate, hard, very hard) and Stimulation (active, sham) as within-subject factors. For the N-back task, a 2 × 2 RM-ANOVA was used with Difficulty (2-back, 3-back) and Stimulation (active, sham) as within-subject factors.

To assess phase-dependent behavioral effects, analogous RM-ANOVAs were conducted using the Fisher z-transformed (pseudo-)r_circ_ values as dependent variables. These analyses tested whether the strength of the phase–behavior relationship differed as a function of stimulation condition and task difficulty. Finally, Rayleigh tests were used to determine whether preferred phases across participants were non-uniformly distributed, indicating clustering around a specific phase angle. Post hoc tests consisted of Bonferroni-corrected paired-samples t-tests. When the assumption of sphericity was violated, Greenhouse–Geisser-corrected values are reported.

## 3. Results

### 3.1. General effects of tACS on working memory

We investigated whether prefrontal theta tACS had a general effect on CDT performance (Figure 2A, 2B). As anticipated, a significant effect of Difficulty was found on the CDT (F(2,38) = 32.37, p < 0.001, η_p_^2^ = 0.63) on accuracy (Figure 2A), where the percentage correct scaled according to the difficulty level (medium: 72.95 ± 1.59%, hard: 64.48 ± 1.65%, very hard: 60.14 ± 1.04% correct). Crucially, however, no significant main effects of Stimulation (F(1,19) = 0.57, p = .459, η_p_^2^ = 0.03) or interaction effect (F(2,38) = 0.41, p = .629, η_p_^2^ = 0.02) were observed. For RT (Figure 2B), no significant main effects of Difficulty (F(2,38) = 0.02, p = .934, η_p_^2^ < 0.01) and Stimulation (F(1,19) = 0.07, p = .792, η_p_^2^ < 0.01), nor an interaction effect (F(2,38) = 0.85, p = .433, η_p_^2^ = 0.04) were found.

**Figure 2.**
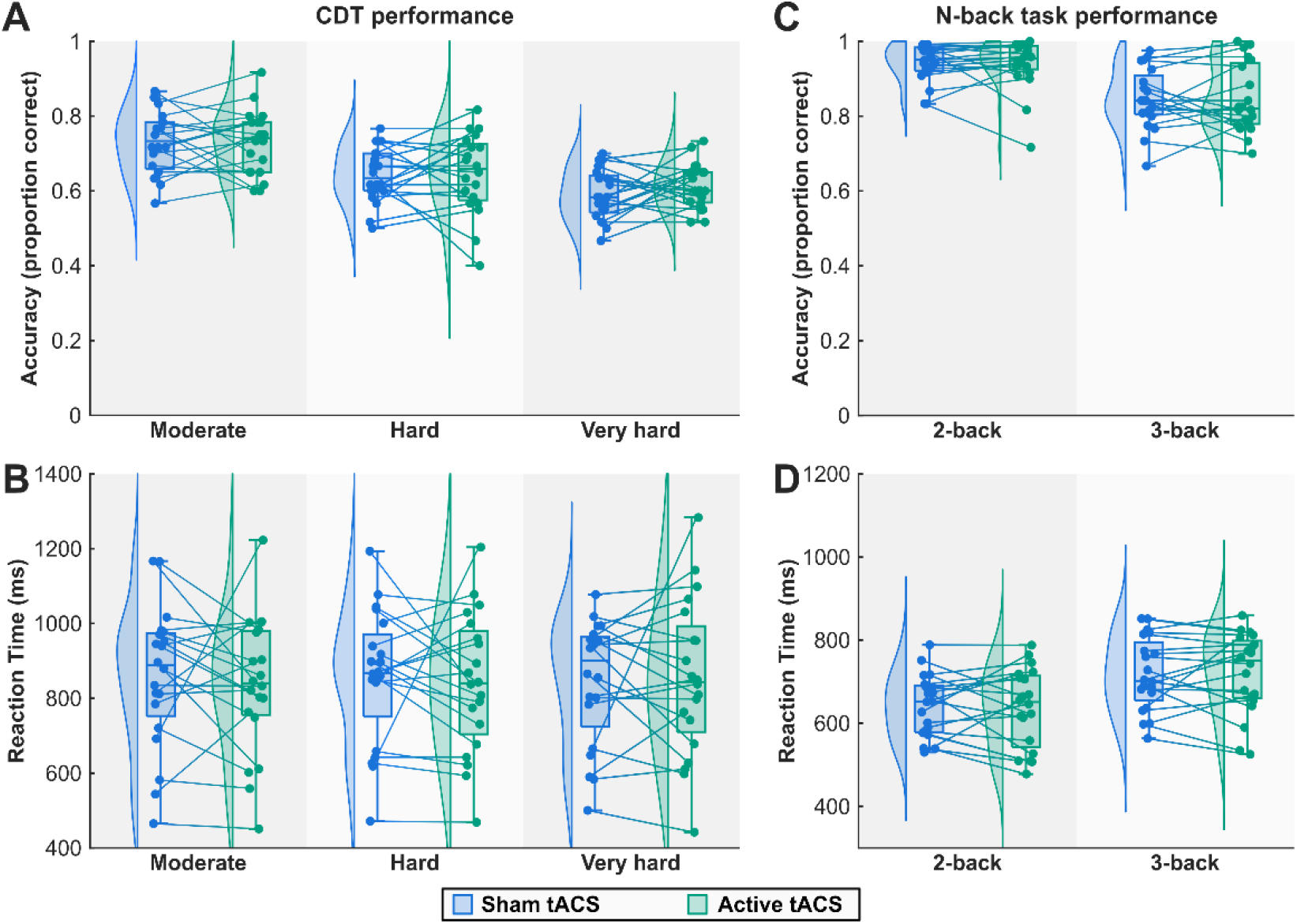
Behavioral results of bilateral prefrontal theta tACS on CDT and the N-back performance. More difficult variations of each task resulted in significantly lower accuracy. Stimulation had no effect on accuracy or RT in either the CDT or the N-back task. A) CDT accuracy, B) CDT RT, C) N-back accuracy, D) N-back RT. Boxplots indicate the median and interquartile range, and density plots show the distribution of values.

A similar analysis was performed on the effects of prefrontal theta tACS on general N-back task performance (Figure 2C, 2D). With accuracy as the outcome measure (Figure 2C), a significant effect of Difficulty was observed (F(1,19) = 48.31, p < 0.001, η_p_^2^ = 0.718), with percentage correct decreasing as difficulty increased (2-back: 94.00 ± 1.16%, 3-back: 84.64 ± 1.76% correct). However, no significant main effect of Stimulation (F(1,19) = 0.01, p = .935, η_p_^2^ < 0.01) or interaction effect (F(1,19) = 0.13, p = .725, η_p_^2^ = 0.01) were found. Similarly, with RT as outcome measure (Figure 2D), a significant effect of Difficulty was found (F(1,19) = 16.61, p < 0.001, η_p_^2^ = 0.47), but not for Stimulation (F(1,19) = 0.01, p = .933, η_p_^2^ < 0.01) nor Difficulty x Stimulation (F(1,19) = 0.24, p = .630, η_p_^2^ = 0.01).

### 3.2. Phase-specific effects of tACS on CDT performance

The analysis with r_circ_ related to RT as the dependent variable (Figure 3A) revealed a significant effect of Stimulation (F(1,19) = 7.11, p = .015, η_p_^2^ = 0.27). On average, phase-locking was higher during active tACS compared to sham (active tACS r_circ_: 0.188 ± 0.011, sham r_circ_: 0.145 ± 0.011). This effect was consistent across the three difficulty levels, as indicated by the absence of a Difficulty x Stimulation interaction effect (F(2,38) = 0.11, p = .874, η_p_^2^ = 0.01). Furthermore, no significant effect on Difficulty was observed (F(2,38) = 1.62, p = .212, η_p_^2^ = 0.08). This finding indicates that the timing of CDT stimuli relative to the tACS cycle predicts the speed of responses in WM.

**Figure 3.**
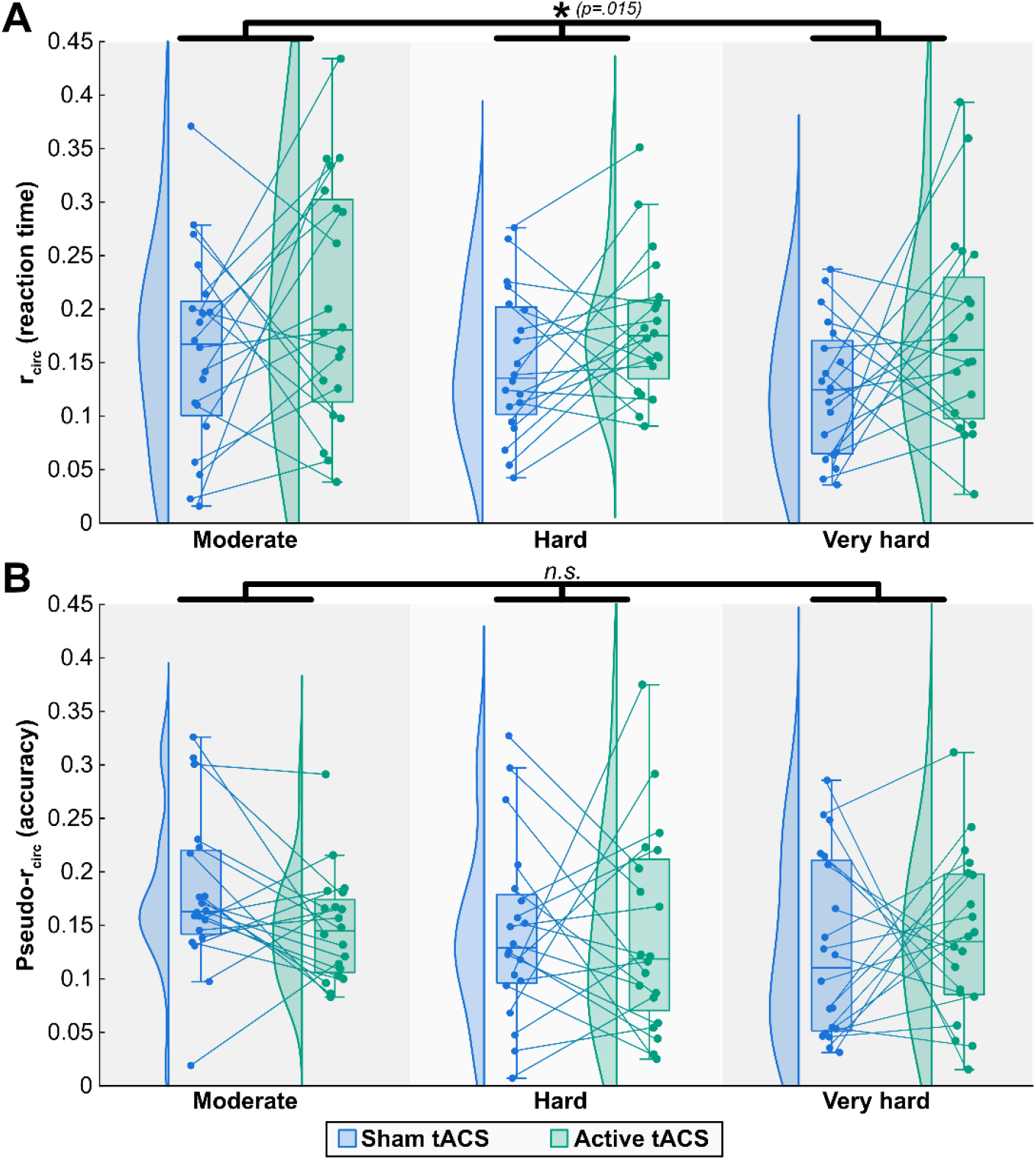
Phase-dependent modulation of RT and accuracy in the CDT. (A) Fisher z-transformed circular–linear correlation coefficients (r_circ_) quantifying the relationship between tACS phase at test-array presentation and RT, shown separately for sham (blue) and active (green) tACS across difficulty levels. (B) Fisher z-transformed pseudo-r_circ_ values quantifying phase-dependent modulation of accuracy, derived from logistic regression models, shown for the same conditions. Points represent individual participants. Boxplots indicate the median and interquartile range, and density plots show the distribution of values. The phase–RT relationship was significantly stronger during active compared with sham tACS, whereas no significant stimulation effect was observed for accuracy. *p<.05; n.s., not significant.

While this result suggests the phase of stimulation affects RT, it is unclear whether such phase locking is consistent across participants. Rayleigh tests were performed per difficulty level during active stimulation (Figure 4), showing no non-uniformity (medium: Z = 0.45, p = .646, hard: Z = 0.80, p = .453, very hard: Z = 0.60, p = .552). Circular-linear regression fits for the highest difficulty level are shown in Figure 5 (same plots for the medium and hard difficulty levels are shown in Supplementary Figures 1 and 2). This observation suggests that while tACS effects are phase-dependent, the preferred phase varies between individuals, with no common phase bias.

**Figure 4.**
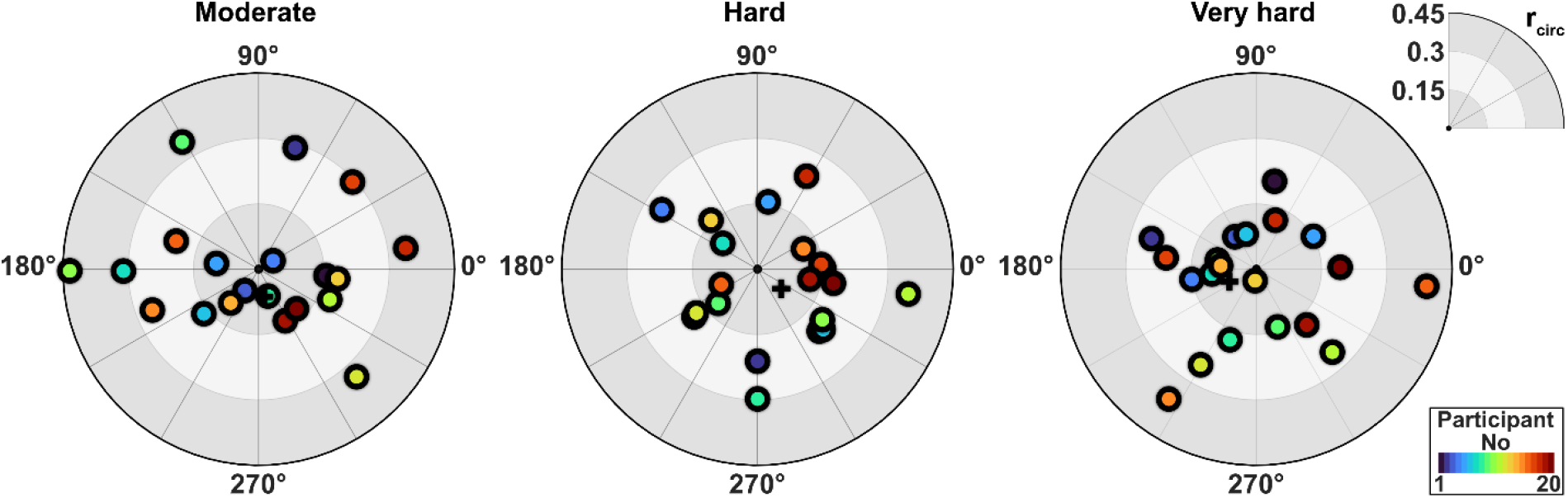
Preferred phase and strength of phase-dependent RT modulation in the change detection task. Polar plots show individual participants’ preferred phases for RT during active tACS, separately for the moderate, hard, and very hard difficulty levels. The angular position indicates the preferred phase, defined as the phase associated with the fastest predicted reaction times. The radial distance from the center reflects the strength of phase-dependent modulation, indexed by Fisher z-transformed r_circ_. Across difficulty levels, preferred phases were distributed across the tACS cycle and did not significantly cluster around a common phase.

**Figure 5.**
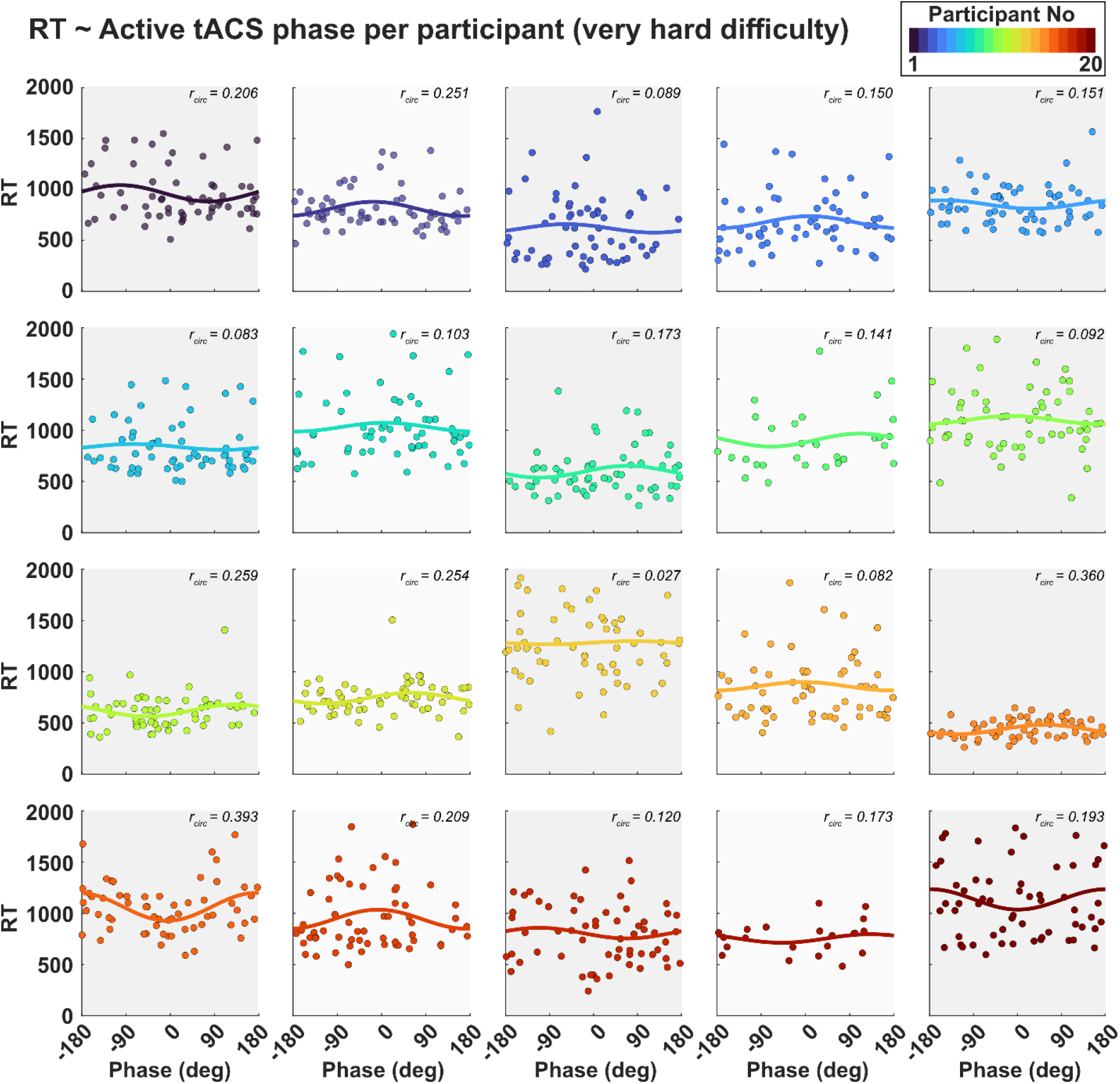
Participant-level phase-dependent RT modulation in the CDT (very hard condition). Individual regression fits show the relationship between active tACS phase at test-array presentation and RT. Each subplot represents one participant. Dots indicate individual trials, and solid lines show the fitted circular regression model. Values in each subplot indicate the strength of phase-dependent modulation, indexed by Fisher z-transformed r_circ_.

The analysis with pseudo-r_circ_ related to accuracy as the dependent variable (Figure 3B) showed no significant effect of Stimulation (F(1,19) = 0.26, p = .616, η_p_^2^ = 0.01). Furthermore, no main effect of Difficulty (F(2,38) = 2.15, p = .140, η_p_^2^ = 0.10) or Difficulty x Stimulation interaction effect (F(2,38) = 0.92, p = .387, η_p_^2^ = 0.05) was observed.

### 3.3. Phase-specific effects of tACS on N-back task performance

The analysis with r_circ_ related to RT as the dependent variable revealed no significant effect of Stimulation (F(1,19) = 0.66, p = .426, η_p_^2^ = 0.03). However, a significant Stimulation x Difficulty interaction effect was observed (F(1,19) = 6.11, p = .023, η_p_^2^ = 0.24). Follow-up paired t-tests suggested that compared to sham r_circ_ values were significantly increased with tACS for the 3-back task (t(19) = 2.50, p = .022, d = 0.56), but not for the 2-back task (t(19) = 1.20, p = .245, d = 0.27). As shown in Figure 6A, phase-locking was higher during active tACS compared to sham in the 3-back task (active tACS r_circ_: 0.297 ± 0.029, sham r_circ_: 0.210 ± 0.029). No significant effect of difficulty was found on phase (F(1,19) = 1.26, p = .275, η_p_^2^ = 0.06). This finding indicates that the timing of 3-back stimuli relative to the tACS cycle predicts the speed of responses in WM.

**Figure 6.**
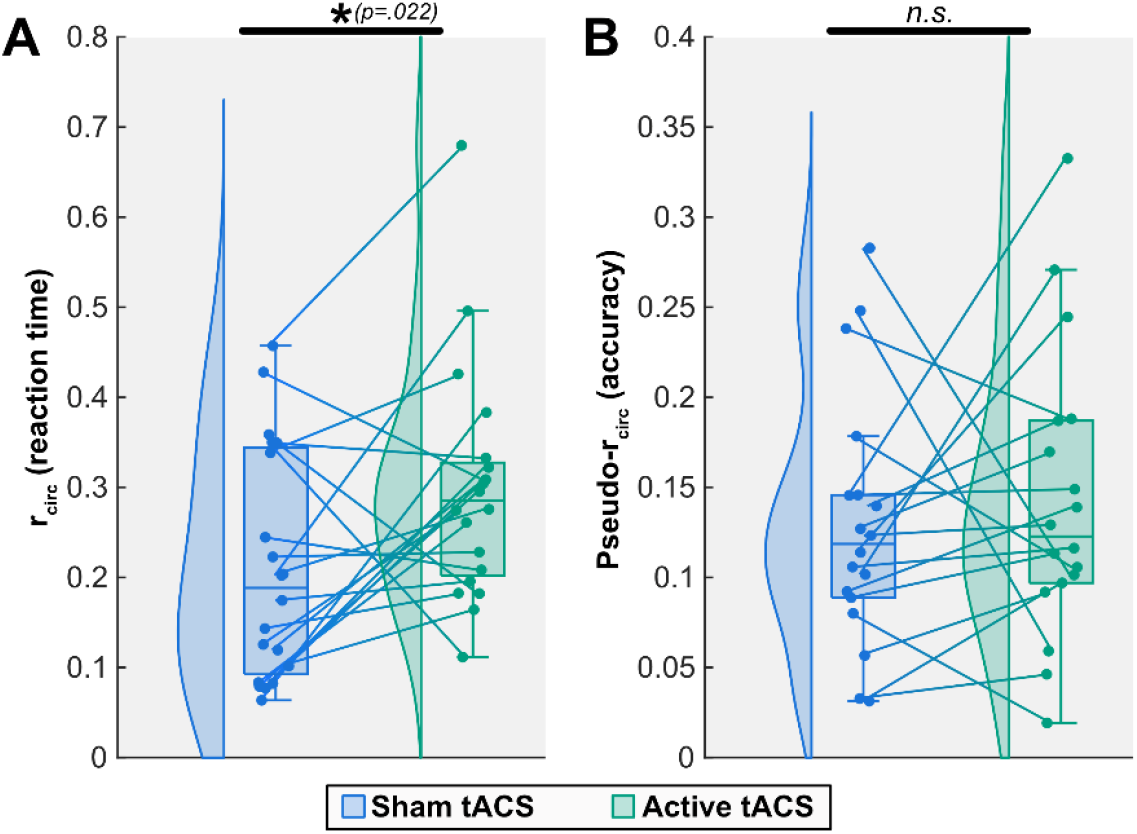
Phase-dependent modulation of RT and accuracy in the 3-back task. (A) Fisher z-transformed circular–linear correlation coefficients (r_circ_) quantifying the relationship between tACS phase at letter presentation and RT, shown for sham and active tACS in the 3-back condition. (B) Fisher z-transformed pseudo-r_circ_ values quantifying phase-dependent modulation of accuracy, derived from logistic regression models, shown for the same condition. Points represent individual participants, with lines connecting sham and active sessions within participants. Boxplots indicate the median and interquartile range, and density plots show the distribution of values. Phase-dependent modulation of RT was significantly stronger during active compared with sham tACS, whereas no significant stimulation effect was observed for accuracy. *p<.05; n.s., not significant.

Subsequently, the consistency of phase locking between participants was investigated. A Rayleigh test was performed for the phase distribution of active tACS in the 3-back task (Figure 7), showing no non-uniformity (Z = 0.34, p = .718). Circular-linear regression fits for all participants are shown in Figure 8. As for the CDT, this observation suggests that while tACS effects are phase-dependent, the preferred phase varies between individuals, with no common phase bias.

**Figure 7.**
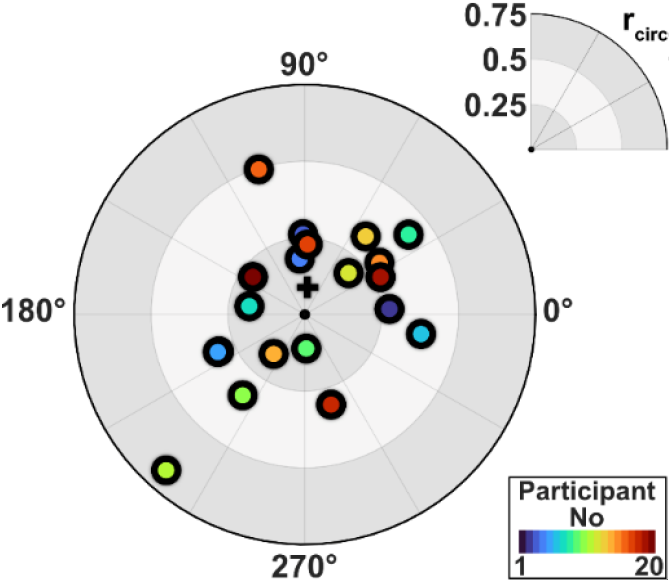
Preferred phase and strength of phase-dependent RT modulation in the 3-back task per participant. The angular position indicates the preferred phase, defined as the phase associated with the fastest predicted reaction times. The radial distance from the center reflects the strength of phase-dependent modulation, indexed by Fisher z-transformed r_circ_. Preferred phases were distributed across the tACS cycle and didnot significantly cluster around a common phase.

**Figure 8.**
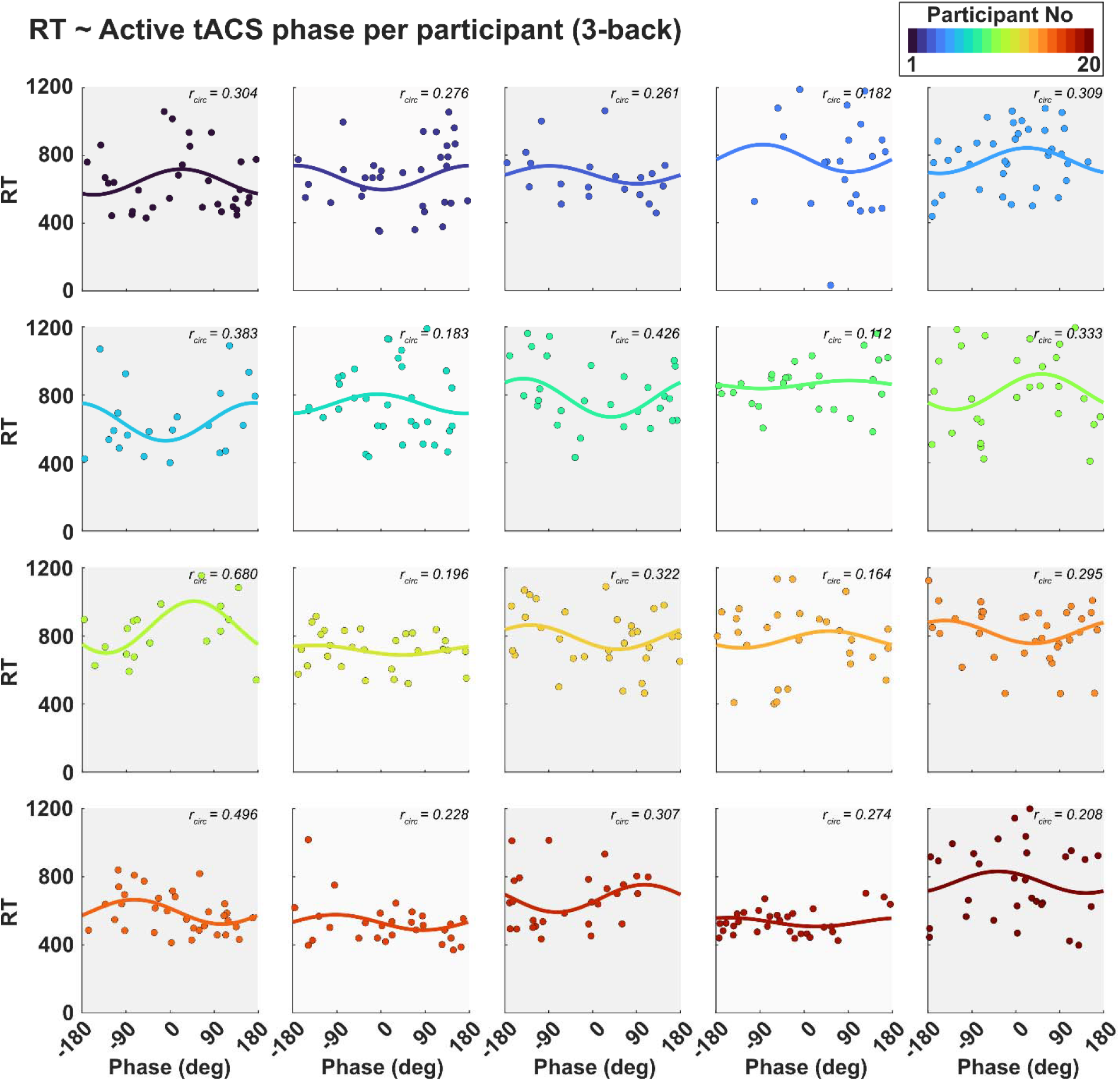
Participant-level phase-dependent RT modulation in the 3-back task. Individual regression fits show the relationship between active tACS phase at test-array presentation and RT. Each subplot represents one participant. Dots indicate individual trials, and solid lines show the fitted circular regression model. Values in each subplot indicate the strength of phase-dependent modulation, indexed by Fisher z-transformed r_circ_.

Phase-locking in relation to accuracy with pseudo-r_circ_ as the dependent variable (Figure 6B) was investigated for the 3-back task. No effect of Stimulation (F(1,17) = 0.29, p = .557, η_p_^2^ = 0.01) was observed. Due to (almost) perfect scores in the 2-back in several participants, no logistic curves could be fitted. As such, no statistical comparison between sham and active stimulation was performed.

## 4. Discussion

By analyzing prefrontal theta-tACS phase and behavioral performance during WM tasks, the present study yielded three main findings. First, overall behavioral performance was not significantly affected by prefrontal theta-tACS, neither for accuracy nor RT in the CDT or N-back task. Second, phase-specific analyses showed that stimulus timing relative to the tACS cycle significantly predicted RT. In the CDT, this effect was observed across all difficulty levels, whereas in the N-back task it was present only in the 3-back condition. Third, preferred phases were not consistently clustered across participants. Together, these findings suggest that prefrontal theta-tACS effects are phase-dependent and show substantial inter-individual variability in the phase associated with optimal performance.

Compared to sham, RTs showed stronger rhythmic modulation when tACS was applied. This suggests that tACS effects are not uniform, but depend on phase alignment between the externally applied oscillation and intrinsic excitability fluctuations. Previous work has shown that stimulation coinciding with phases of high neural excitability facilitates neural responses, whereas stimulation at phases of lower excitability may reduce responses [40,41,43]. Besides physiological responses, tACS phase also affects behavioral performance, such as the detection of visual stimuli [42,61]. In line with our results, De Graaf et al. [61] found that the tACS phase specifically affects RTs in target detection. While their results stem from a different task, region, and frequency, it supports the idea that tACS phase may affect response speeds in behavioral tasks. Here, this effect was observed in both verbal and spatial WM, suggesting that it is not limited to a single domain. Together, this may indicate that tACS phase influences the temporal gating of information processing. As such, a tentative hypothesis is that information arriving at a non-preferred phase is processed with a brief delay, resulting in slower RTs, whereas information arriving at the preferred phase can be processed more directly, resulting in faster RTs. Indeed, this interpretation aligns with electrophysiological work suggesting that neural oscillation phase acts as a temporal gating mechanism [62,63].

While a phase-dependent RT effects were observed at the participant level, the preferred phase (i.e., the phase associated with the fastest RTs) varied across individuals, and no generalizable preferred phase emerged. Several factors may explain this variability. First, the same externally applied tACS phase may not produce identical cortical effects across individuals. Individual differences in cortical folding and tissue geometry can alter how the electric field reaches neural tissue, meaning that the scalp-defined stimulation phase may map differently onto cortical excitability across participants [64–66]. Second, inter-individual differences in endogenous theta frequency may have contributed to variability in preferred phase. Here, a fixed 4 Hz stimulation frequency was used to systematically assess phase-dependent effects under identical stimulation conditions. Because endogenous theta frequency varies across individuals, the same tACS phase may correspond to different intrinsic oscillatory states across participants, which may explain why no consistent preferred phase emerged at the group level. Third, we defined phase relative to the time at which WM stimuli were presented. Naturally, stimulus information is not available to the prefrontal cortex instantaneously, but reaches prefrontal WM networks through bottom-up processing pathways [67]. Individual differences in the speed of these processes may introduce variability between stimulus-locked tACS phase and the phase at which information actually reaches prefrontal cortex [68], further contributing to the observed variability in preferred phase.

In contrast to the phase-dependent effects observed for RT, we found no evidence that tACS phase predicted task accuracy. This absence of phase-dependent accuracy effects can be interpreted in two ways. First, as discussed above, tACS phase may primarily affect processing speed, but not accuracy. Second, our stimulation parameters may not have been optimal for modulating task accuracy. For instance, the tACS montage used in the present study (F3, F4) is commonly used in WM research [28,34,56,69]. This setup affects dorsolateral and dorsomedial prefrontal regions, while recently it was reported that targeting the ventrolateral and ventromedial prefrontal cortex may be more effective at modulating WM performance [30]. As such, it cannot be ruled out that a phase-dependent effect would have been found using a different montage. Previously, it was shown that the phase of alpha oscillation in the visual cortex is associated with the accuracy of visual detection, both in electroencephalography and tACS studies [42,70,71]. As such, the absence of phase-dependent effects on accuracy in this study should be interpreted with caution and warrants further research.

Several limitations should be considered. First, stimulation was delivered at a fixed 4 Hz frequency across participants. This standardized phase assessment, but leaves open whether individualized theta-frequency stimulation would yield stronger or more stable effects. Second, no concurrent neural recordings were obtained. During tACS, the stimulation artifact overlaps with the frequency of interest, making endogenous theta phase difficult to extract reliably. Although artifact-removal approaches exist [72–74], they mainly recover oscillatory power and may not preserve the original neural phase. Therefore, our analyses relate behavior to the applied stimulation phase rather than directly measured endogenous theta activity. Third, phase-dependent N-back effects were observed only in the 3-back condition. Because 2-back accuracy was very high, this condition may have been insufficiently demanding to reveal phase-dependent modulation and should therefore be interpreted cautiously.

In conclusion, the efficacy of theta-tACS during working memory may depend not only on where or at which frequency stimulation is applied, but also on when task-relevant information is presented relative to the stimulation cycle. Although prefrontal theta-tACS produced no overall behavioral benefit, phase-specific analyses revealed that RTs depended on stimulus timing within the stimulation cycle. Thus, disregarding stimulation phase may obscure behaviorally relevant tACS effects. Moreover, the optimal phase differed across individuals, suggesting that phase may be a hidden source of variability in tACS studies. While personalized stimulation approaches have focused on optimizing location and frequency [75], phase alignment between stimulation and task events may represent an additional dimension of personalization.

## CRediT authorship contribution statement

**Julie Dimmendaal:** Conceptualization, Formal analysis, Investigation, Writing - Original Draft. **Xiaojuan Wang:** Conceptualization, Writing - Original Draft. **Bas J. Dijkslag:** Investigation, Writing - Review & Editing. **Laura E. Huizinga:** Investigation, Writing - Review & Editing. **Sander Maalderink:** Investigation, Writing - Review & Editing. **Marijn Priest:** Investigation, Writing - Review & Editing. **Femke J.E. van Dam:** Investigation, Writing - Review & Editing. **Mark M. Span:** Methodology, Software, Resources, Writing - Review & Editing. **Miles Wischnewski:** Conceptualization, Methodology, Formal analysis, Writing - Original Draft, Visualization, Supervision, Funding acquisition.

## Data availability

Data and analysis code are available on Zenodo: [76].

