## Supplementary Data for "Phase of transcranial alternating current stimulation modulates working memory processing speed"

**Supplementary materials**

Julie Dimmendaal^1^, Xiaojuan Wang^1,2^, Bas J. Dijkslag ^1^, Laura E. Huizinga^1^, Sander Maalderink^1^, Marijn Priest^1^, Femke J.E. van Dam^1^, Mark M. Span^1^, Miles Wischnewski^1,3,^*

^1^Department of Psychology, University of Groningen, Groningen, The Netherlands

^2^Medical School, Tianjin University, Tianjin, China

^3^Department of Cognitive Neuroscience, Maastricht University, Maastricht, The Netherlands

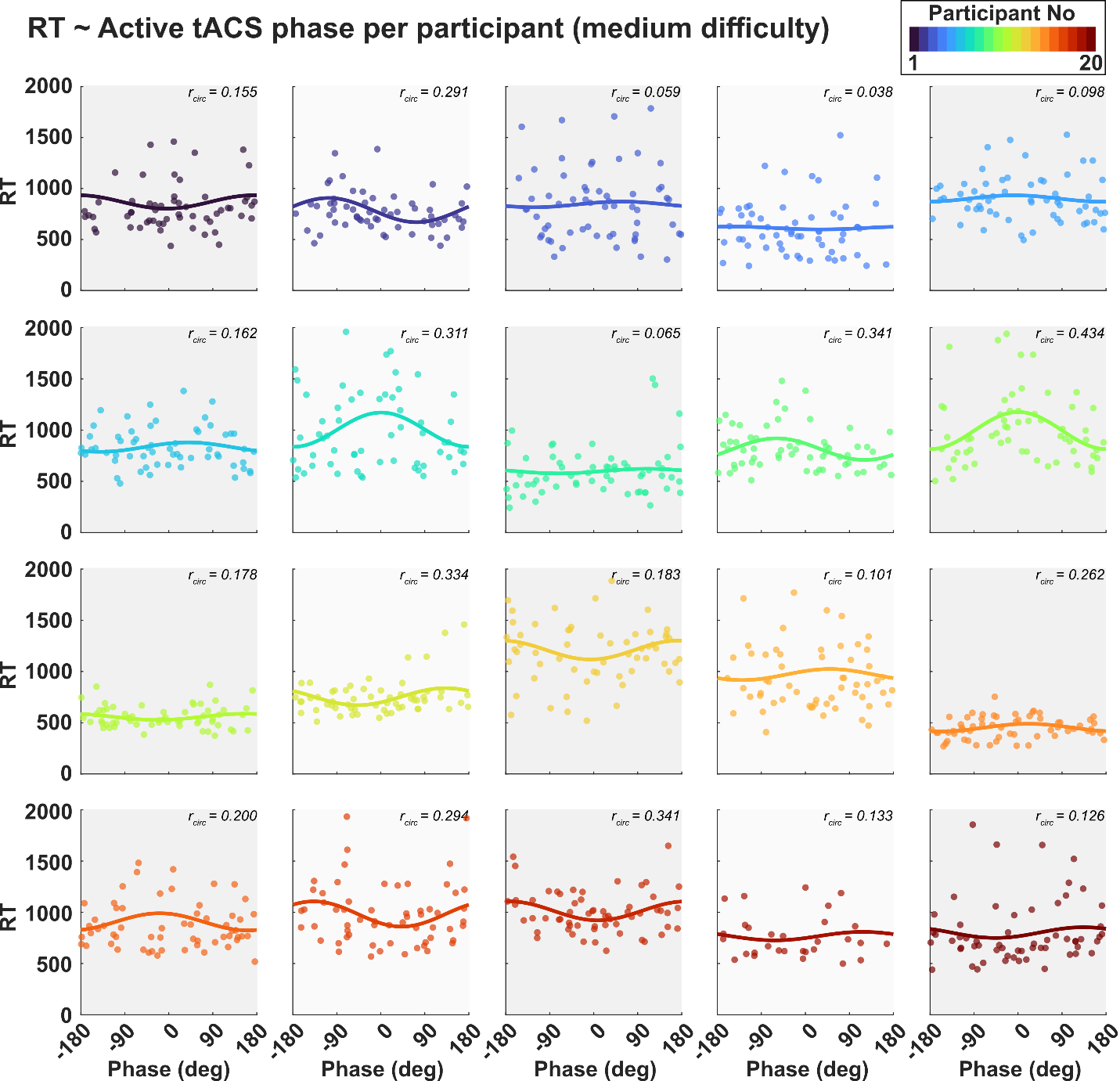
**Supplementary Figure 1.** Participant-level phase-dependent RT modulation in the CDT (medium condition). Individual regression fits show the relationship between active tACS phase at test-array presentation and RT. Each subplot represents one participant. Dots indicate individual trials, and solid lines show the fitted circular regression model. Values in each subplot indicate the strength of phase-dependent modulation, indexed by Fisher z-transformed r_circ_.

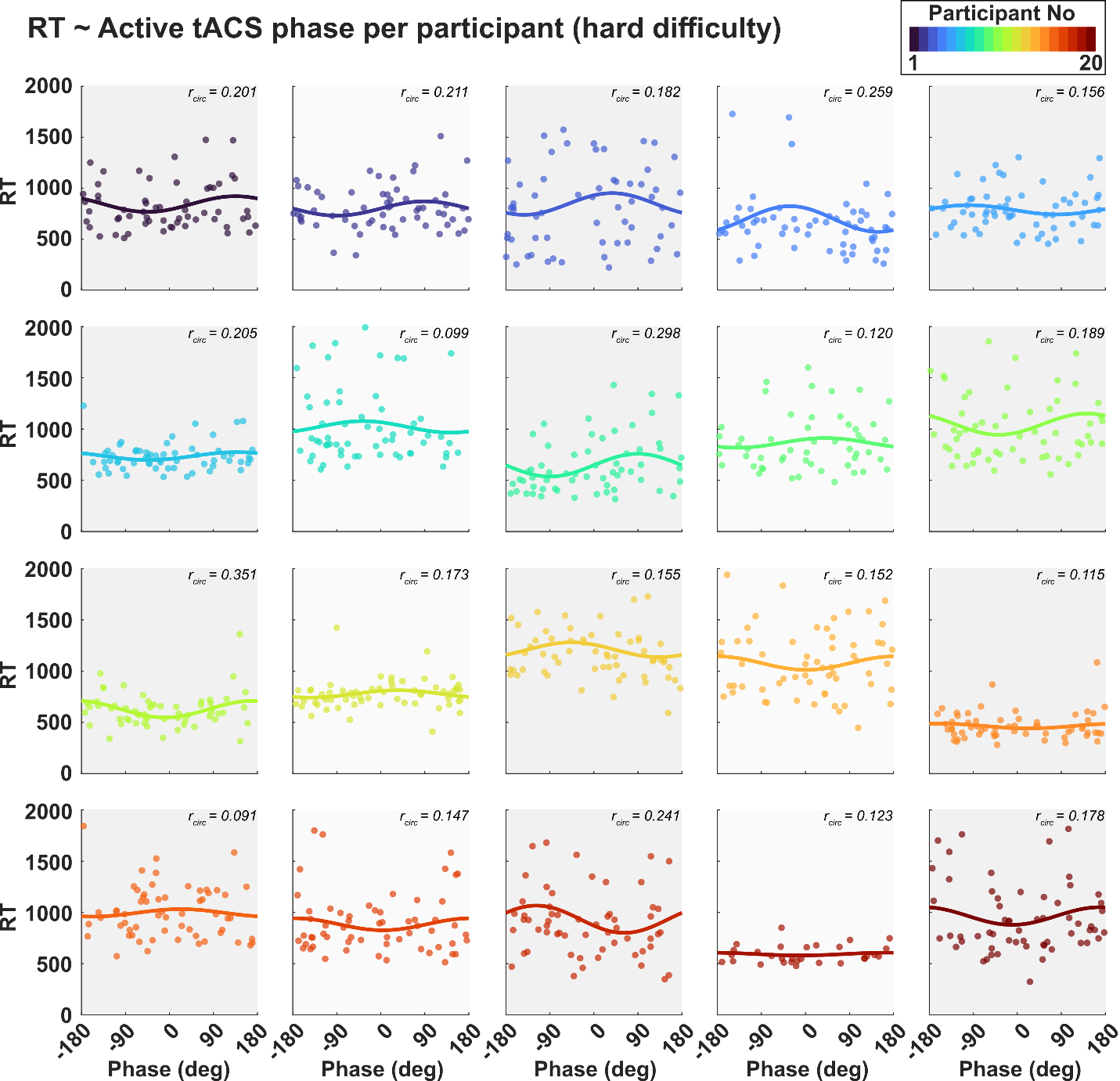

**Supplementary Figure 2.** Participant-level phase-dependent RT modulation in the CDT (hard condition). Individual regression fits show the relationship between active tACS phase at test-array presentation and RT. Each subplot represents one participant. Dots indicate individual trials, and solid lines show the fitted circular regression model. Values in each subplot indicate the strength of phase-dependent modulation, indexed by Fisher z-transformed r_circ_.
